# Spatial mapping of human lung tissue reveals granuloma diversity and histopathological superstructure in tuberculosis

**DOI:** 10.1101/2022.07.31.502240

**Authors:** Andrew Sawyer, Ellis Patrick, Jarem Edwards, James S Wilmott, Timothy Fielder, Qianting Yang, Daniel L Barber, Joel D Ernst, Warwick J Britton, Umaimainthan Palendira, Xinchun Chen, Carl G Feng

**Affiliations:** School of Medical Sciences, Faculty of Medicine and Health, The University of Sydney, Sydney, NSW, Australia; Centenary Institute, The University of Sydney, Sydney, NSW, Australia; Charles Perkins Centre, The University of Sydney, Sydney, NSW, Australia; School of Mathematics and Statistics, Faculty of Science, The University of Sydney, Sydney, NSW, Australia; Centre for Cancer Research, Westmead Institute for Medical Research, The University of Sydney, Westmead, NSW, Australia; Sydney Precision Data Science Centre, The University of Sydney, Sydney, NSW, Australia; Melanoma Institute Australia, The University of Sydney, Sydney, NSW, Australia; Department of Tissue Pathology and Diagnostic Oncology, Royal Prince Alfred Hospital, Camperdown, NSW, Australia; Guangdong Key Lab for Diagnosis & Treatment of Emerging Infectious Diseases, Shenzhen, Third People’s Hospital, Shenzhen, Shenzhen, Guangdong, China; Laboratory of Parasitic Diseases, National Institute of Allergy and Infectious Diseases, National Institutes of Health, Bethesda MD USA; Division of Experimental Medicine, Department of Medicine, University of California, San Francisco CA USA; Department of Clinical Immunology, Royal Prince Alfred Hospital, Camperdown, NSW Australia; Guangdong Key Laboratory of Regional Immunity and Diseases, Department of Pathogen Biology, Shenzhen University School of Medicine, Shenzhen, Guangdong, China; The University of Sydney Institute for Infectious Diseases, The University of Sydney, Sydney, NSW, Australia

**Author notes:** Corresponding authors: Carl Feng or Xinchun Chen.

**Keywords:** tuberculosis, granuloma, lung, human, spatial

## Abstract

The histopathological hallmark of tuberculosis (TB) is the formation of immune cell enriched aggregates called granulomas, but the scope of granuloma heterogeneity in human TB is unknown. By spatially mapping individual immune cells across large regions of TB lung tissue, we report that in addition to necrotizing granulomas, the human TB lung contains abundant non-necrotizing leukocyte aggregates surrounding areas of necrotizing tissue. These cellular lesions were more diverse in composition than necrotizing lesions and could be stratified into four general classes based on cellular composition and spatial distribution of B cells and macrophages, indicating there are foci of distinct immune reactions adjacent to necrotizing granulomas. We further show that the specific cellular composition of non-necrotizing structures correlates with their proximity to necrotizing lesions. Together, our study shows that during tuberculosis diseased lung tissue develops a histopathological superstructure comprising at least four different types of non-necrotizing cellular aggregates organized as satellites of necrotizing granulomas.

## Introduction

Tuberculosis (TB) remains a major cause of mortality with 1.5 million deaths in 2020^1^. Development of new TB vaccines and therapeutics remains a global priority and will depend on a clearer understanding of mechanism mediating host resistance to *Mycobacterium tuberculosis* (M.tb), the causative agent of TB^2–4^. The battle between the host and the pathogen occurs at the level of immune cell-enriched lesions called granulomas^5^. While being recognized as a protective structure for walling-off infection, granulomas also contribute to the survival of mycobacteria and importantly are the major manifestation of organ damage^6–9^.

Despite the realization of the critical role of granuloma in host defence, the cellular basis of mycobacterial granuloma formation is incompletely defined. Investigations of granulomas in human tissues are hampered by the lack of access to clinical samples and limited imaging technologies. In addition, commonly used laboratory mouse strains do not form lesions with the key features of human granulomas^10–12^ These hurdles are further compounded by the challenges in granuloma classification due to the high variation in lesion size and histopathological features in diseased human tissues^13,14^. Consequently, most studies choose to characterize one or two types of lesions (eg, non-necrotizing and necrotizing) in isolation. The extent of variation across non-necrotizing TB lesions is unknown.

Recent advancement in spatial transcriptomics and proteomics technologies enable researchers to image mycobacterium-infected tissues at various levels. In this regard, spatial transcriptomics have provided comprehensive gene expression information across tissue sections of human leprosy^15^ and murine TB^16^, although the current technologies do not have the resolution required for single-cell imaging. Antibody-based multiplexed tissue imaging approaches^17,18^, including one with multiplexed ion beam imaging by time of flight (MIBI-TOF), have been employed to examine human TB granulomas with single-cell resolution. The MIBI-TOF study has successfully revealed diverse micro-environments within human TB granulomas^18^. However, this approach interrogates mainly the selected regions of individual lesions, its scope in tissue-wide cross-lesion comparative analysis is limited ^19,20^. Because only a subset of lesions were analyzed, it is unclear whether the observed difference in microenvironments reflects the cellular structural variations within a lesion or across multiple lesions. Therefore, revealing the level of granuloma diversity by analyzing many lesions across large areas of diseased tissue will deepen the understanding of the intra-lesion microenvironment characterized by high-dimension phenotyping.

To capture all cellular structures present in a TB lung tissue sample, we have employed multiplex Opal immunofluorescence to map the tissue-wide immune landscape (size ranging from 100 to 300 mm^2^, ten TB patients in total) with single cell resolution. Moreover, we developed a novel granuloma identification and classification strategy to minimize the subjectivity and bias commonly associated with lesion analysis. By performing single-cell spatial analysis on 657 lesions (defined as distinct immune cell aggregates) with approximately 7 million cells, we established that human TB lung tissues contain, in addition to well-recognized necrotizing lesions, at least four types of compositionally and spatially distinct non-necrotizing lesions. Further spatial analysis demonstrated that the latter lesion structures are shaped by the neighbouring necrotizing granuloma as well as by intra-lesion organization of immune cells, suggesting that granuloma architecture is governed by lesion-intrinsic and -extrinsic signals in human TB.

## Results

### Active pulmonary TB is associated with extensive tissue consolidation and development of immune cell aggregates in the lungs

To gain information on tissue-wide inflammatory response to M.tb infection in human lungs, we first examined H&E-stained tissue sections obtained from ten patients who underwent partial surgical lung resection due to unsuccessful TB drug therapy. Major histopathological changes were evaluated by two independent histopathologists. We observed that in contrast to healthy lungs with clear alveolar space, the landscape of diseased tissues was highly complex with regions of consolidation and foci of leukocytes (Fig. 1A). The well-recognized necrotizing granulomas were frequently accompanied by many non-necrotizing granulomas and lymphoid aggregates. The superstructure was consistently observed across all patient samples (n=10) (Fig. 1A and 1B), suggesting that the multifaceted histopathological landscape is common in the lungs of active TB patients.

**Figure 1.**
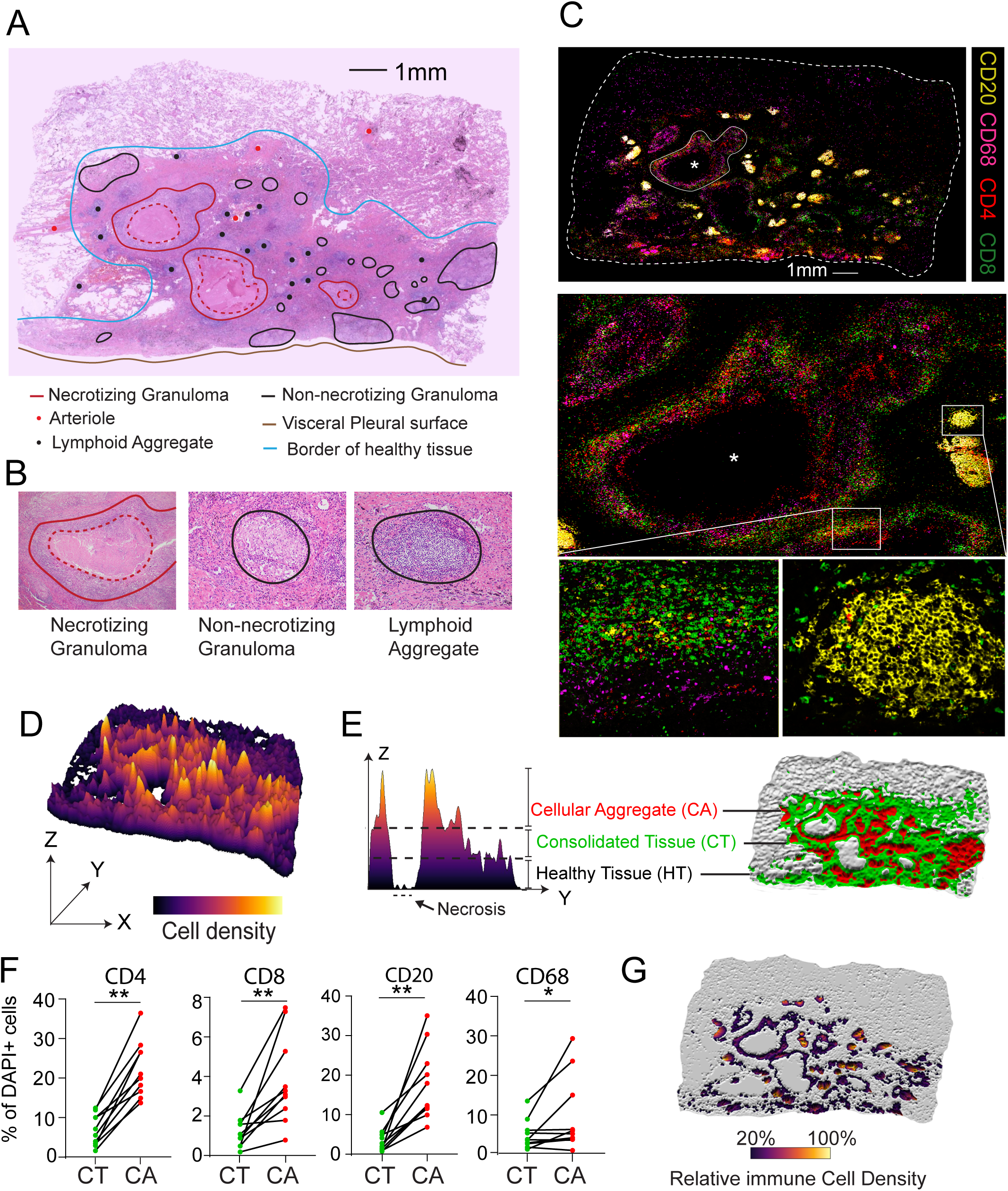
Active pulmonary TB is characterized by development of histopathological superstructures in the lung. **(A)** A TB lung tissue section stained with H&E with various histological and histopathological features indicated. **(B)** Enlarged example images of distinct lesion types identified in (A). The outer and inner boarders of necrotizing lesions were marked with solid and dotted red lines, respectively. The boarders of non-necrotizing lesions were marked with solid black lines. **(C)** Composite Opal immunofluorescence image of the same tissue section in A stained for major immune populations inlaid with magnified views of a necrotizing lesion (top panel, circled by a white line, *necrotizing core) and two non-necrotizing regions (middle panel, rectangle boxes). **(D)** 3D diagram showing the variations in the density of DAPI^+^ segmented cells across entire TB lung section from (A). The height of the peak (Z-axis) indicates cell density. **(E)** A representative 2D histogram showing a cross-section (indicated by the white dotted line in C) along Y- and Z-axis. The three Z-axis segments (divided by dotted lines) are defined based on cell density and correspond to the regions of healthy, consolidated tissue and cellular aggregates shown in the pseudo colour topography image (right) of the same TB lung tissue as in A and C. **(F)** Percentage of major immune cell populations among total DAPI positive cells in consolidated tissue (CT, green) versus cellular aggregates region (CA, red). Each paired point represents a patient. The level of statistical significance is determined using a Wilcoxon signed-rank test and a p-value < 0.05 was considered statistically significant. * *P* < 0.05 ** *P* < 0.01. **(G)** Distribution of immune cell-containing cell aggregates (coloured) across the same tissue section as in (A). The lesions selected are high in cellularity and positive for at least one of four major immune cell lineage markers (CD4, CD20, CD8 or CD68). Colour intensity of non-greyed areas correlates reversely with immune cell density of the region relative to the maximum density in the tissue.

To visualize the immune landscape across entire lung sections, tissues were stained with the major immune lineage markers CD68 (Macrophages), CD20 (B cells), CD8 (CD8^+^ T cells) and CD4 (CD4^+^ T cells), together with nuclear dye DAPI (Fig. 1C). High-resolution 20X regional images acquired using the Akoya Vectra system were stitched digitally to reconstruct the images of entire plane of the tissue section, allowing the visualization of each tissue section in its entirety and at a single cell resolution. We observed extensive infiltration of immune cells across all tissue sections, which was mirrored in the HE-stained serial sections from the same patient. In addition, giant cells were observed across patients and proliferative (Ki-67+) cell aggregates were observed in two patients (Fig. S1A and B).

To assess whether the diverse structures defined pathologically could be captured and analyzed at single-cell level, we first examined nuclear dye DAPI expression on TB lung tissue sections. To objectively distinguish healthy lung tissue from consolidated areas and cellular aggregates in the immunofluorescence images, healthy and inflamed regions were first stratified based on the relative density of nucleated cells on the tissue-wide Opal IHC image following cell segmentation with DAPI using HALO software (Indica Labs) (Fig. S1C). Kernel density estimation was employed to quantify the cell density. The variations in the density across TB lung tissue were visualized with 3D projection (Fig. 1D), where height of the peaks (y-axis) indicates cell density. Each distinct peak represents a cell aggregate, and a trough (coloured in white) reports an acellular necrotizing lesion. Tissue was subsequently divided into regions based on cell density relative to the densest area of each patient sample. Cellular aggregates were identified as areas with density between 50-100% of the maximum density, consolidated tissues between 25-50% and healthy lung tissue as below 25%. When mapped back onto the tissue, the localization of the cell density-defined, healthy tissue (HT) consolidated tissue (CT) and cellular aggregates (CA) were matched closely to the regions identified on HE-stained sections (Fig. 1E).

Next, we compared the composition of major immune cell populations in the consolidated region and cellular aggregates and found that, as a proportion of total DAPI^+^ cells, CD68^+^, CD8^+^, CD20^+^ and CD4^+^ cells were all significantly more abundant in the cellular aggregates compared to the consolidated region (Fig. 1F). These observations indicate that these immune cell-enriched cellular aggregates, defined by our unbiased cell density-based method are distinct lesions within the consolidated region and represent the “granulomas” and lymphoid aggregates defined conventionally with histopathological features. To capture the total immune cell aggregates more precisely, we refined our lesion identification strategy by performing kernel density analysis and lesion definition only on cells stained positively for immune markers (Fig. 1G). In the reminder of this investigation, we have employed this unbiased analytic method to define individual immune cell aggregates in human tissues. We have focused our analysis on cellular aggregates containing 20-100% of the maximum immune cell density per patient, which included 672 lesions across ten patients which included 22 structures containing a central necrotizing cavity. Together, this unbiased lesion identification approach has revealed previously unappreciated complexity in the cellular landscape of human TB granulomas.

### Cellular aggregates are highly diverse and fall across a continuous spectrum underpinned by the density of CD20^+^ cells

Opal IHC images have revealed that the immune cell compositions in TB lesions are highly diverse (Fig. 2A). To understand the lesion heterogeneity, we analysed the density of each immune cell type across 672 lesions. For necrotizing lesions, the central area of necrosis was excluded from the density calculation to allow meaningful comparison with non-necrotizing lesions. We observed that the density of individual immune cell population varied significantly across lesions (up to 3 logs) and interestingly the variation was more apparent in non-necrotizing lesions than their necrotizing counterparts (Fig. 2B). CD20^+^ B cells and CD4^+^ T cells were more abundant in non-necrotizing lesions while CD68^+^ macrophages were more abundant in necrotizing lesions. CD8^+^ T cells distributed indistinguishably between necrotizing and non-necrotizing lesions.

**Figure 2.**
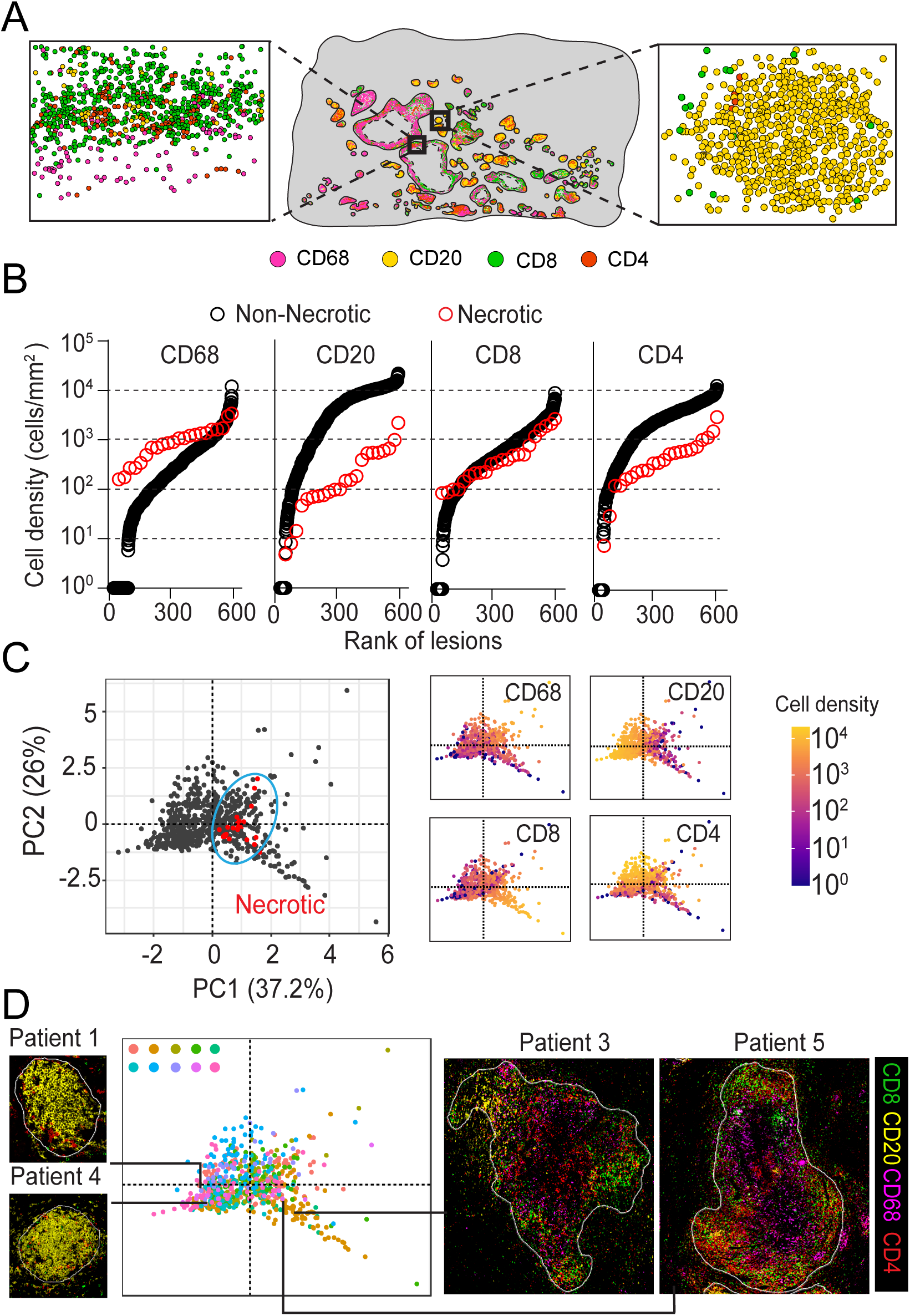
Lesions have diverse compositions but fall across a continuous spectrum underpinned by CD20^+^ cell density. **(A)** Image of segmented immune cells in positively identified lesions in TB lung from Fig. 1A with two representative areas (boxed) showing distinct cellular composition. **(B)** Necrotizing and non-necrotizing lesions from 10 patients ranked according to the density of each major immune cell populations. Each symbol represents an individual lesion. **(C)** PCA plot of all lesions (n=657) with necrotizing lesions indicated in red and circled by an ellipse. Additional marker-specific PCA plots are coloured according to density of marker positive cells. **(D)** PCA plot visualization of all lesions across 10 patients. Each colour indicates one patient, and each symbol represents an individual lesion. Images of example lesions of four individual patients are shown.

To assess whether the lesions cluster into distinct groups based on their immune cell composition, we performed Principal Component Analysis (PCA) analysis on 672 lesions from ten patients based on the density of four major immune cell populations. Interestingly, we did not observe clustering of lesions into distinct groups, although necrotizing lesions (colored in red) tend to cluster closely (Fig. 2C), indicating a continuous spectrum of lesion compositions. PC1 represented the largest share of variation (37.2%) and was primarily driven by the variation in the density of B cells, indicating the continuous spectrum of the lesions is underlined mainly by the variations in the CD20^+^ cell density. Moreover, these lesions did not cluster according to patient with the exception of one individual patient with increased density of CD8^+^ T cell (Fig. 2D), suggesting a minimal patient variation in lesion types defined using 4 immune cell markers.

### Lesions can be stratified based on immune cell composition and spatial organization

Having established a spectrum of lesion compositions based on immune cell density, we sought to determine whether lesions have diverse spatial arrangement of cells. As expected, necrotizing lesions were characterized by their acellular central core (Fig. 3A). However, within non-necrotizing solid lesions, cells were also found to distribute unevenly, in some lesions the majority of the cells preferentially localized in the central region whilst in others most cells localized peripherally (Fig. 3A) leaving a loose but non-necrotizing central area. To analyze this spatial variation, we devised a method termed “total cell Central Preference Index (tCPI)” to give a single number readout of whether a lesion tends to have an increased or reduced density in its central area compared to the average density of the lesion (Fig. 3B and 3C). Briefly, the metric works by measuring the distance of all cells from the lesion’s outer border and then dividing cells into an inner 50% of cells and outer 50% of cells. The area covered by the outer 50% of cells is divided by the total lesion area and will give an output of less than 0.5 if there is reduced density of cells in the central area of the lesion. As expected, when the tCPI was compared between necrotizing and non-necrotizing lesions, the former lesions had significantly lower central preference index than non-necrotizing lesions (Fig. 3D), validating the approach in quantifying cell spatial distribution tendency in mycobacterial granulomas.

**Figure 3.**
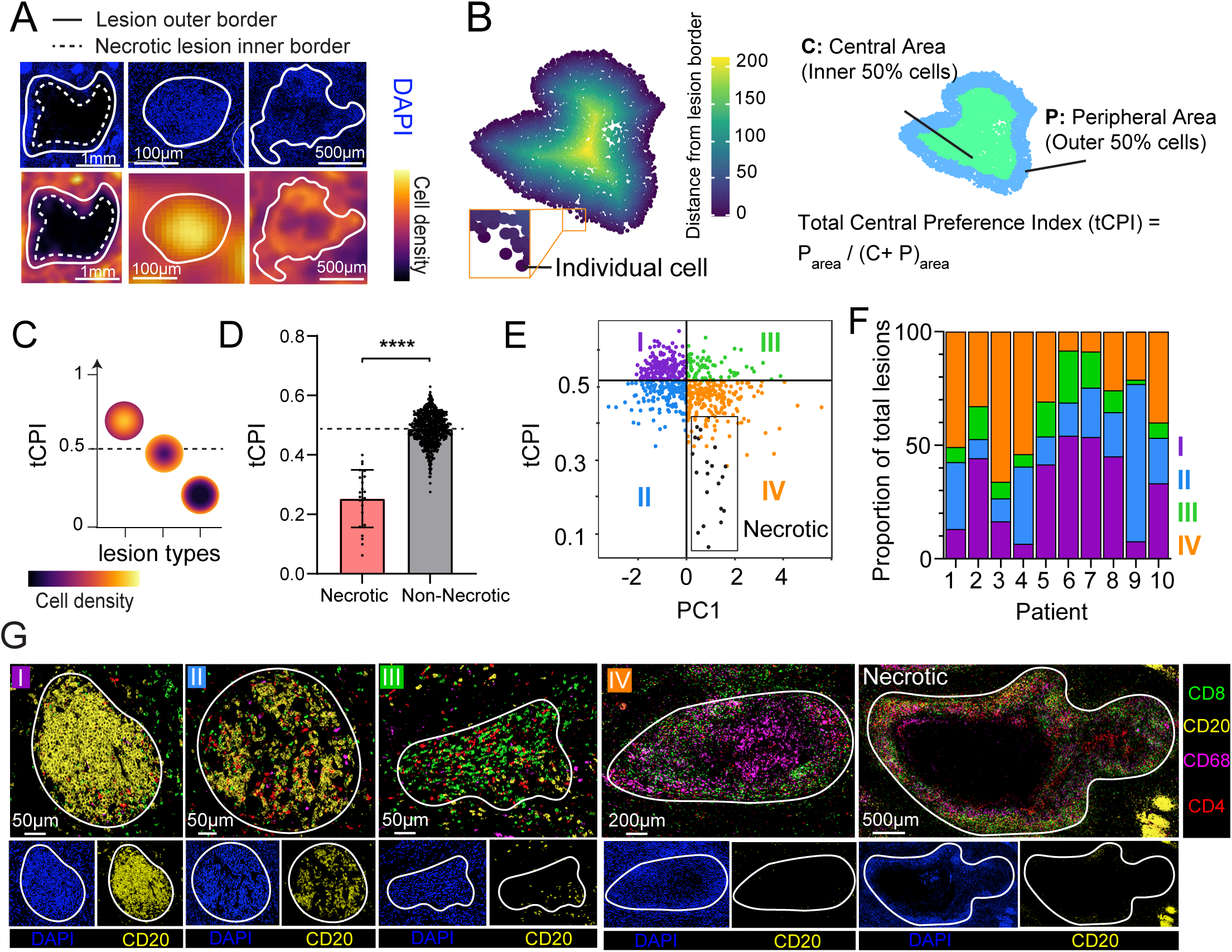
Combined analysis of compositional and spatial features identifies four types of TB lesions. **(A)** Images of DAPI-stained tissue showing variations in intra-lesion cell distribution (top panel). Heatmap images showing the same three lesions with relative cell density indicated (Lower panel). **(B)** A lesion image with the distance of each individual cell from the lesion boarder indicated by colour density (Left). The same lesion with areas coloured according to which cells are closer or further from the lesion border relative to the median distance (Right). Equation used to calculate total cell central preference index is shown. **(C)** Schematic diagram illustrating the intra-lesion cell distribution patterns and their corresponding tCPI. **(D)** tCPI compared between necrotizing and non-necrotizing lesions. Each symbol represents an individual lesion. The level of statistical significance is determined using a Mann-Whitney test, **** *P* < 0.001. **(E)** Pairwise comparison of PC1 from Fig. 2C against tCPI. Each symbol represents an individual lesion. Lesions (n=657) are divided into 4 sub-populations with the quadrants being set at PC1 = 0 and tCPI = 0.5. The 5^th^ type of lesions, necrotizing lesions (black symbols), are additionally indicated in the boxed area. **(F)** Proportion of each lesion type across 10 patients. Each color represents one lesion type. **(G)** Representative multiplex images of each lesion type with correlated images of DAPI and CD20 staining shown below each multiplex image.

We next investigated if the information of cell density and the tendencies of immune cells to locate toward or away from the lesion centre could facilitate lesion stratification. We performed pairwise comparison of PC1 (driven mainly by B cells) from the previous PCA (Fig. 2C) and central preference index measurement. We termed this unbiased lesion identification and classification algorithm LANDSCAPE (Lesion And Neighbourhood Description Stratified by cellular Composition And Position Estimates). In addition to necrotizing lesions readily identifiable on H&E-stained sections, we used this analysis to define four types of solid lesions(Fig. 3E). Interestingly, all 4 lesion types were identified in each of the 10 patients analyzed although there were variations in the proportion of each lesion type across patients (Fig. 3F, Fig S1G), suggesting that the lesion types identified here represent common immune features of the local tissue response in human TB lungs.

The cellular characteristics of the lesion type defined using LANDSCAPE analysis were highly consistent with those presented on multiplexed immunofluorescence images of the same lesion type (Fig. 3E and 3G). Type I lesions frequently have more B lymphocytes and cells tend to localize close to the lesion centre (high tCPI) (Fig. S1D, S1E and S1F). Type II lesions also have a higher numbers of B cells but with reduced cell density in the lesion centre (low tCPI). Both type III and IV lesions had reduced presence of B cells and increased macrophages. However, they exhibited distinct cellular organization. Type IV lesions displayed lower tCPI than type III lesions, suggesting that in these lesions most cells localized preferentially away from lesion centres. This description is reinforced by the observation that all necrotizing / cavitary lesions fell into type IV region (lesions shown in the box area).

### Divergent intra-lesion spatial distribution of CD20^+^ cells and CD68^+^ cells determine the types of non-necrotizing granulomas

To understand more specifically how the spatial distribution of individual immune cell populations regulates granuloma formation, we quantified the relative distance of lymphoid cells and macrophages from the lesion border and termed the measurement as “immune cell Central Preference Index (immCPI) (Fig. 4A, B). This measurement functioned independently of the density of respective cell-types in each lesion (Fig. S2A) and CPI (Fig. S2B). We found that CD20^+^ cells were located more peripherally in type IV lesions compared to types I, II and III, while CD68^+^ cells were located more centrally in type III and IV compared to type I and type IV compared to type II (Fig. 4C). Together, these data highlight further cellular structural differences between lesion types and suggest a progression of CD20 cells and CD68 cells between lesion types.

**Figure 4.**
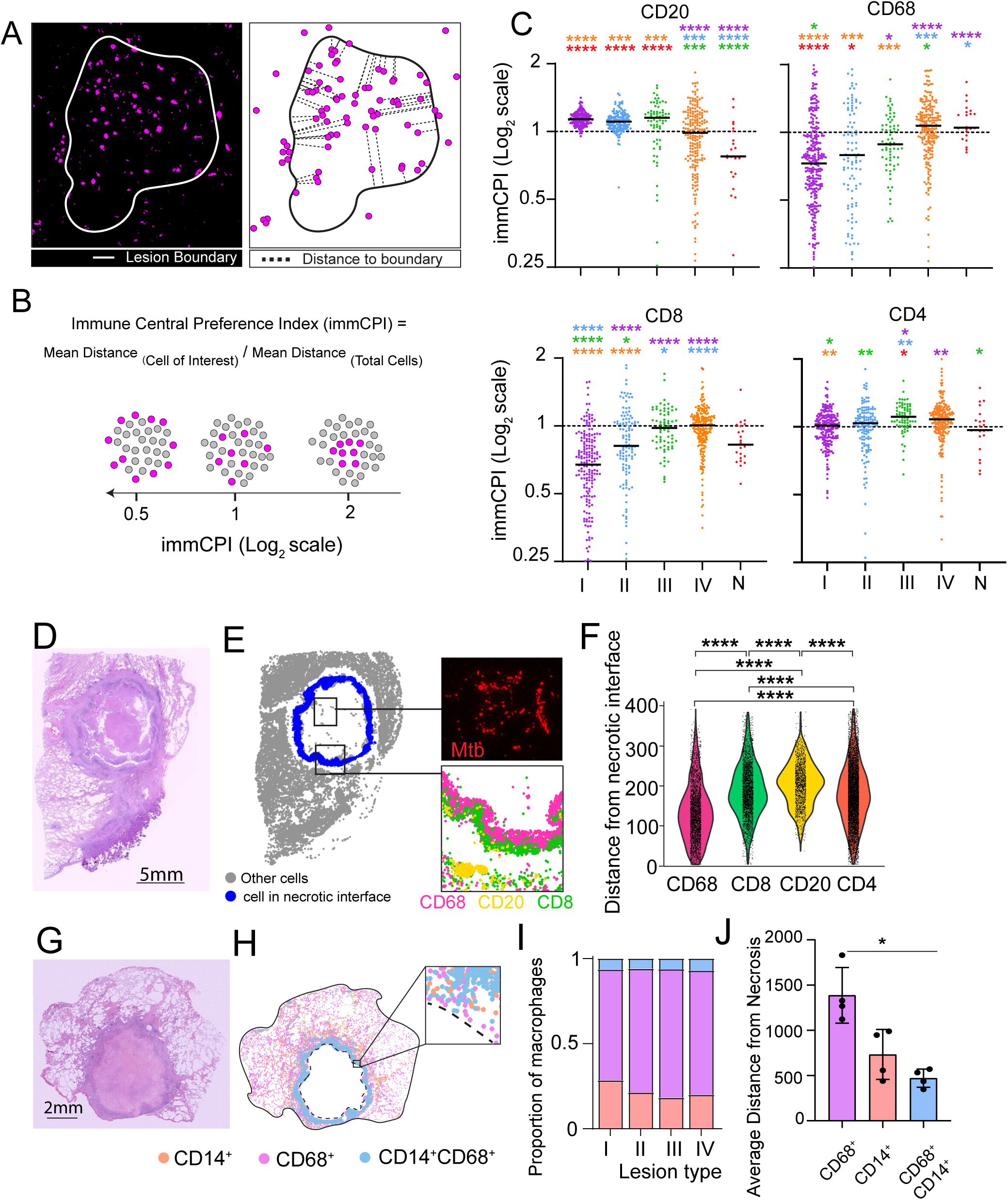
Spatial organization of CD68^+^ macrophages is lesion type-dependent. **(A)** A lesion image with CD68 staining (Left) and a diagram illustrating measurement of the closest distance of individual CD68^+^ cells from the lesion boarder (Right). **(B)** Formula for calculating the spatial distribution cell-of-interest within a lesion. Schematic diagram of the lesions with representative immCPI measurements. **(C)** immCPI of each immune cell type across lesion type. Each symbol represents an individual lesion. **(D)** Image of a H&E-stained TB lung sample. **(E)** Pseudo colour image of the section from (D) with the necrotizing boarder indicated in blue. The magnified images of the boxed areas show positive M.tb staining and spatial position of CD68^+^, CD20^+^ and CD8^+^ cells. **(F)** Distance of individual immune cell population from necrotizing border in a representative lesion. Each symbol represents an individual cell. **(G)** image of a H&E-stained TB lung sample. **(H)** Spatial distribution of monocyte / macrophage populations defined by CD14 and CD68 in tissue section from (G) with magnified image showing individual segmented myeloid cells. **(I)** Proportion of each macrophage subset by lesion type across the tissue section from G. **(J)** Distance of each macrophage subset from closest necrotizing area across entire tissue section. Each symbol represents an individual patient and bars indicate standard deviation. The level of statistical significance in panel C, F and J is determined using a Kruskal-Wallis Test with Dunns Multiple Comparisons Test and an adjusted p-value less than 0.05 was considered statistically significant. * *P* < 0.05, ** *P* < 0.01, *** *P* < 0.001, **** *P* < 0.0001.

### CD14^+^CD68^+^ subset of monocyte / macrophage populations preferentially localize at the necrotizing boarder

Necrotizing granulomas are formed when a host immune system fails to contain M.tb replication or regulate the local inflammatory response ^21^. The pathological manifestation is frequently associated with active disease, tissue injury, and transmission of M.tb. However, there is little knowledge as to how the infection niche is walled of and how immune cells are spatially organized at the boarder of necrotizing (acellular) regions. We co-stained TB lung section with antibodies specific to immune cells and mycobacteria to determine the localization of infection foci in lung sections. We found that M.tb was only positively identified in a subset of tissue sections, almost exclusively in the cavity region (Fig. 4D and E), which is consistent with previous histopathological studies ^22–24^. Curiously, necrotizing lesions also had a higher proportion of KI-67^+^ replicating CD4^+^ T cells compared to other non-necrotizing lesion types (Fig. S2C). To map the cellular microenvironment of necrotizing border, we measured the distance of each immune cell in necrotizing lesions from the necrotizing interface and found that CD68^+^ cells were localized the closest to necrotizing core, followed by CD8^+^ and CD20^+^ cells (Fig. 4F, Fig. S2D).

We next stained a subset of patient samples (n = 5) with an additional myeloid marker CD14 to better understand the populations of macrophages/monocytes residing in TB lung tissues (Fig. 4G and 4H). The relative abundance of the three subtypes, CD14^+^ C68^—^, CD14^—^ CD68^+^ and CD14^+^CD68^+^, was comparable across all four types of granulomas (Fig. 4I). However, while at the whole tissue level, CD68^+^CD14^+^ macrophage/monocyte population was the least abundant subset relative to other two populations, they localized significantly closer to the necrotizing core (Fig. 4J), revealing previously unrecognized tissue-wide spatial distribution of different myeloid cell populations from the necrotizing interface. Future studies focusing on the phenotypes and functions of these three macrophage/monocyte populations in necrotizing lesions may reveal mechanisms underlying infection progression in TB.

### The distance to necrotizing lesions correlates with the relative abundance of specific immune cell populations in the surrounding non-necrotizing lesions

Our tissue-wide imaging analysis revealed that necrotizing lesions are surrounded by diverse solid lesions (Fig. 1 and Fig. 5A), which raises the questions of the spatial relationship between necrotizing and various non-necrotizing lesions. To determine the inter-lesion relationship, we measured the minimum distance from each lesion to every other lesion on the same tissue section (Fig. 5B). Given the cellular organisation we observed around necrosis, we compared the distance of lesions of each type to the closest necrotizing lesion. We found that lesions of type I and II tend to be twice as close to a necrotizing lesion as lesions of type III and IV (Fig. 5C).

**Figure 5.**
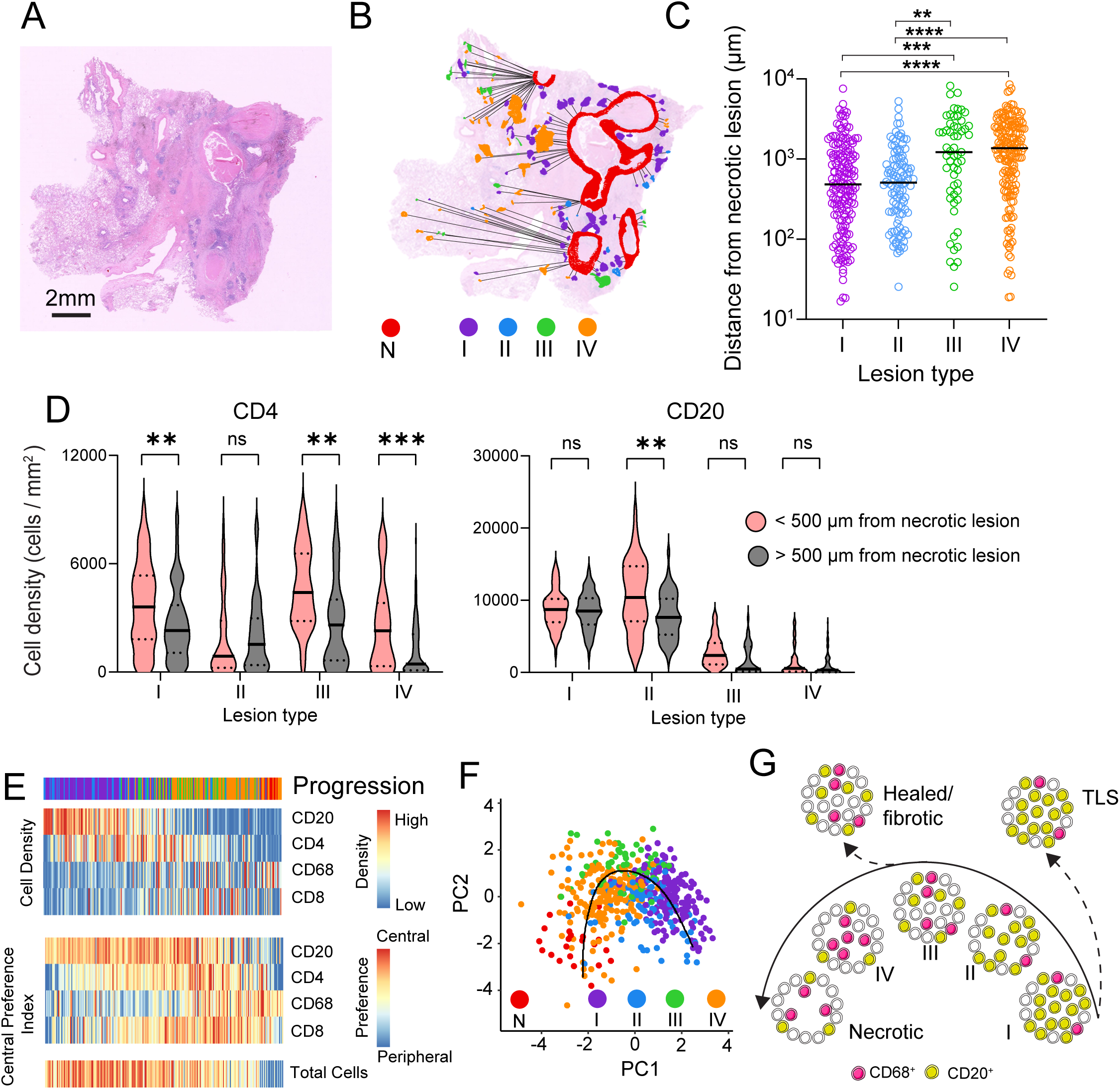
Formation of non-necrotizing lesions is shaped by the neighbouring necrotizing granuloma and intra-lesion organization of CD20^+^ and CD68^+^ cells. **(A)** Image of a H&E-stained TB lung sample. **(B)** Same TB lung section as in (A) overlaid with the locations of non-necrotizing and necrotizing lesions identified using LANDSCAPE tool. The relationship of each non-necrotizing lesion to closest necrotizing lesion is indicated with connecting lines. **(C)** Distance of 4 types of non-necrotizing lesions to the closest necrotizing granuloma. Each symbol represents an individual non-necrotizing lesion. The level of statistical significance is determined using a Kruskal-Wallis Test with Dunns Multiple Comparisons Test and an adjusted p-value less than 0.05 was considered statistically significant. ** *P* < 0.01, *** *P* < 0.001, **** *P* < 0.0001. **(D)** Density of CD4^+^ and CD20^+^ cells in lesions located greater or less than 500 µm from necrotizing lesions. The level of statistical significance is determined using a Mann-Whitney test and a p-value less than 0.05 was considered statistically significant. ** *P* < 0.01 and *** *P* < 0.001. **(E)** Heatmap of inter-lesion trajectory analysis based on spatial and compositional parameters of individual lesions defined by LANDSCAPE. Lesions are ranked along the X axis according to trajectory and manually ordered on the Y axis. Top panel indicates lesion type. The analysis was performed using R. **(F)** Trajectory analysis of all lesions based on spatial and compositional lesion parameters as shown in (E). Line indicates inferred trajectory. Each symbol represents an individual lesion (n=657). **(G)** Schematic diagram summarizing key findings and illustrating hypothetical lesion progression pathways. Dotted lines indicate hypothetical lesion progression branchpoints and pathways.

To further explore the role of non-necrotizing lesion distance from necrotizing lesions, we divided lesions of each type into those localized greater than 500 µm from necrosis and those less than 500 µm. We found that the lesions of type I, III and IV had more CD4^+^ cells when they localized closer to necrotizing lesions (< 500 µm) compared to those further away from necrosis (< 500 µm) (Fig. 5D). Type II lesions contain more CD20^+^ cells when they were closer than 500 µm compared to when it was further. There was no difference in the density of CD8^+^ or CD68^+^ cells in any lesion type when comparing the distance of the lesions from the closest necrotizing lesion (Fig. S2E). These results suggest that potential cross-talks between necrotizing and certain types of non-necrotizing lesions may contribute to the formation of multi-lesion superstructures in the lung.

### Formation of non-necrotizing lesions is shaped by the proximity to necrotizing granulomas and intra-lesion organization of CD20^+^ and CD68^+^ cells

Since granulomas are highly dynamic and diverse, lesion-lesion interactions may also play a role in regulating lesion progression and the overall lung pathology. We combined both compositional and spatial measurements taken thus far and performed trajectory analysis on all lesions. We found that lesion progression correlates with the order of the four types of lesions identified and that necrotizing lesions consistently fall at one end of spectrum (Fig. 5E and 5F). CD20^+^ cell density, as well as the central preference of total leukocytes, CD20^+^ and CD68^+^ cells all follow along the progression trajectory closely. Together, we hypothesize that that the lesions identified here may represent distinct yet related stages of granuloma maturation, from B cell-enriched lymphocytic foci to macrophage-centred granulomas, and some of the latter lesions may eventually develop into necrotizing tissue whereas others become healed or fibrotic lesions. Similarly, type I lesions may contain both distinct tertiary lymphoid structures (TLS) and B cell-enriched infection foci (Fig. 5G). Future high-dimension phenotyping of the four lesion types on samples with different clinical outcomes will assist in defining the branchpoints along the lesion progression.

## Discussion

Recent work in M.tb-infected non-human primates and patients has revealed that TB lesions are heterogenous in their histopathological features^17,25,26^, metabolic activities^27,28^, inflammatory signalling^29,30^ and pathogen burdens^9,31,32^. Collectively, these previous studies have contributed to the notion that TB represents a spectrum of clinical and sub-clinical manifestations^12,13,33^. By mapping immune landscape at a single cell level across entire lung tissue section, we demonstrate here that even on the same lung section, human TB lesions, especially non-necrotizing cellular aggregates, are highly diverse in their composition and cellular spatial organization. Furthermore, these non-necrotizing lesions, together with necrotizing granulomas, appear to form a histopathological superstructure underlying the immunopathology associated with uncontrolled M.tb infection. Since each lesion type represents a distinct type of immune response and lesions may individually or cooperatively contribute to TB pathogenesis, we suggest that both intra- and inter-lesion analysis should be performed to define host response to M.tb.

By analyzing > 650 lesions with a total of 7 million single cells, we defined four types of non-necrotizing lesions representing a spectrum of distinct yet overlapping cellular architecture. The frequently documented lesions with a necrotizing core represent only one of the several types of lesions in the infected organs, suggesting that examination of only one or two stereotypical lesion types is unlikely to capture all the cellular events contributing to the overall lung pathology and clinical outcome. It is of particular interest that each type of the four different solid lesions was present in all the patients examined. We speculate that the lesion spectrum may reflect the history of ongoing local immune response to continuing seeding of locally disseminated drug-resistant bacteria. Therefore, this heterogenous and dynamic nature captured in a single tissue section may be explored to identify distinct cellular structures associated with many if not all of the stages of the granuloma maturation process. A similar approach has been used to successfully define T cell population at different maturation stages in the thymus.

We have revealed the extent of granuloma heterogeneity in patients with ongoing M.tb infection. Interestingly, PCA analysis showed that TB granulomas are organized as a spectrum rather than distinct clusters, suggesting overlaps in cellular structure and potentially interactions across lesions. The spatial association between certain types of non-necrotizing and necrotizing granulomas further suggests that these lesions may function cooperatively as an inflammatory superstructure to determine the outcome of local immune response. These observations raise a question how a TB granuloma should be defined at a single cell level. It is impossible to determining with 2-dimentional immunofluorescence staining whether these structures were structurally connected as has been shown reported previously with μCT imaging^26^. Nevertheless, our results support the hypothesis that the “double-edged sword” function of granulomas may reflect the heterogeneity among granuloma populations^33^.

Advances in single-cell RNA sequencing have provided unprecedented insights into the composition and function of immune cells in M.tb-infected tissues^34–37^. However, this approach does not provide the spatial context of the local immune response^38^, which is critical in understanding the architecture and function of mycobacterial granulomas. By employing MIBI-TOF, McCaffrey *et al*. have successfully documented high-dimension cellular information of human granulomas^18^. However, though time-of-flight mass cytometry can be used to analyse the regions of interest within a lesion, it does not reveal comprehensive tissue-level immune landscape or lesion-lesion relationship^19^. On the other hand, our tissue-wide single-cell analysis is better suited for examining the tissue landscape across lesions at the cost of detailed phenotyping of immune populations. Therefore, the two studies are highly complementary. Comparing and contrasting the data generated in the two investigations may provide additional insights into human TB granuloma structure. For example, some of our B cell-enriched type I lesions, which may resemble tertiary lymphoid structures (TLS) formed in mycobacterial infected tissues^39^, have also been independently identified in the McCaffrey study^18^. Therefore, it is possible that Type I lesions represent heterogenous lesions including conventional TLS and B cell-enriched mycobacterial granulomas. Future deep-phenotyping using high-dimensional image technology will be able to address this question.

Our study has several limitations. Firstly, a limited number of lung tissue samples were examined. While a minimal patient variation was observed when four immune cell markers were used, the variation between patients may become more apparent when more markers are used to profile a broader variety of immune subsets. Secondly, Opal multiplex technology is only capable of examining small numbers of markers thereby limiting the depth of cellular analysis. Combining tissue landscape and in-depth region of interest analysis will lead to a better understating of lesion microenvironment in the context of surrounding inflamed tissues. Finally, since our initial samples were collected from TB patients undergoing lobectomy due to a poor treatment response, whether our study has captured a fraction or full spectrum of granulomatous lesions remains to be determined. In future, we will apply our LANDSCAPE strategy to broaden our investigation by including samples from patients with active TB or latent infection in lungs resected for other reasons, which will assist in elucidating the relationship between cellular structure, anti-microbial function and disease outcomes.

## Materials and Methods

### Patient Cohort

We utilized a retrospective study cohort of patients from the Shenzhen Third People’s Hospital (Shenzhen, China). The study was approved by the Research Ethics Committee of the Shenzhen Third People’s Hospital (protocol ID #: 2016-081). All archival clinical specimens were analyzed, with no active participation of human subjects. Active TB diagnosis was based on clinical symptoms, chest radiography, pathological result, microscopy for acid-fast bacilli, M.tb culture, and GeneXpert analysis of sputum (Tab. S1). Serial sections (5 μm) of each specimen were stained with H&E and inspected by two clinical pathologists to evaluate the presence of granulomatous inflammation. Patients with comorbid HIV were excluded from the study. All patients with TB were undergoing anti-TB therapy.

### OPAL staining

Formalin-fixed paraffin-embedded (FFPE) TB lung sections were baked at 65°C for 1 hour and dewaxed and rehydrated through gradients of Xylene and ethanol concentrations. Antigen retrieval was performed by boiling samples to 100°C for 15 minutes in pH 9 Antigen retrieval buffer (Akoya). Tissues were incubated for 45 minutes at room temperature with primary antibodies specific for CD20 (L26, Cell Marque, 1:1000), CD4 (SP35, Cell Marque, 1:100), CD68 (Kp-1, Cell Marque, 1:500), CD8 (ab4055, Abcam, 1:500), CD14 (HPA001887, Cell Marque, 1:100), Ki-67 (D2H10, Cell Signalling technologies, 1:1000) or M.tb (Rabbit polyclonal Ab to M.tb, Cat# ab905, Abcam). Primary antibodies were detected with OPAL Polymer HRP Ms + Rb (Akoya Biosciences) for 15 minutes and visualized following a 10-minute incubation with Tyramide signal amplification and Opal Fluorophore (Akoya) before being boiled at 100°C to strip antibody-HRP complexes from tissue. The process was repeated until all markers have been stained. Following the final marker staining, tissue section was incubated with DAPI for 10 minutes and mounted using ProLong Gold (Thermofisher). The images were captured using the Vectra 3.0.5 multispectral imaging system (Akoya Biosciences). A series of high-power ×20 multispectral images (MSIs) covering entirety of tissue sections were captured.

### Cell Segmentation

MSIs were unmixed using Inform 2.3 (Akoya) and exported as .qtiff images. Tiff images were then imported in HALO3.3.2541.345 (Indica Labs) and digitally stitched to become construct single images of entire tissue sections. HALO was trained to segment individual cells based on characteristic nuclear staining in the DAPI channel. To distinguish cell identities the positivity threshold for each marker was determined and cell segmentation was performed. Object Data (including identity and location information for each cell) for of each patient sample was exported and further analysis was performed in R x64 3.5.3^40^

### Density and Immune-cell marker-based lesion identification

Kernel density analysis was performed using the “`kde2d`” function in the MASS^41^ package in R using bandwidth of 500 within a 500 by 500 grid. 3D projections were created with the “`persp`” function and coloured with the “`viridis`” function from the Viridis package^42^. Relative to the maximum cell density in each patient the healthy region was defined by densities between 0-25%, consolidated region between 25-50% and aggregate region between 50-100%. To visualize the regions defined by cellular density on tissue images the Density Heatmap tool in HALO was used applied to map density across entire tissue section and the Annotation tool was used to define the regions for export.

To refine the lesion identification, Kernel Density Analysis was further performed on lesions that were stained positively for at least one of four immune cell markers (CD20, CD68 CD4 or CD8) in R. 3D density projection was again performed with “`persp`” function and areas between 25-100% of the maximum cell density were considered as diseased tissue areas. Lesions were identified using HALO with the density heatmap function and lesion boundaries were drawn using the annotation tool. Cell segmentation was performed in batch on all lesions and object data was exported to R.

### Lesion clustering analysis based on intra-lesion immune cell density

The area size of individual lesion was determined using HALO summary data in R. The relative abundance of individual immune cell populations in each lesion was expressed as immune cells per mm^2^ lesion area. Lesion clustering analysis was performed using PCA with density of CD8^+^, CD20^+^, CD4^+^ and CD68^+^ cells used as parameters. PCA was performed in R with the “`prcomp`” function with “`scale = True`”. PCA results were visualised using the factoextra package^43^ with functions “`fviz_pca_ind`” and “`fviz_pca_var`”.

### Total cell central preference index, Immune cell central preference index and Necrotic distance index calculation

The Infiltration Analysis tool in HALO was applied to each lesion to measure the distance of each cell from the lesion border, with store object data set as true to save individual cell data. Data were imported into R and cell distance information was matched to previous cell data by cell ID. To determine the Central Preference Index of all DAPI^+^ cells (tCPI), median cell distance from the border was identified with the median function in R. The area of lesion encompassing the inner/outer 50% of cells was then calculated in HALO using the infiltration tool. The tCPI was calculated as:

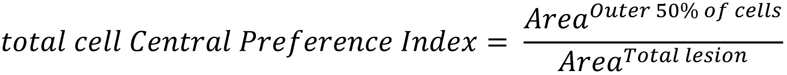

For Immune Cell Preference Index (immCPI) determination, cell distance measurements from lesion borders used in tCPI calculation were applied to cells based on their immune cell lineage identity. Relative distance of each marker from the lesion border was calculated as follows:

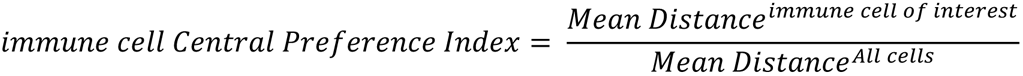

Relative distance of each marker from the necrotizing interface was calculated as follows

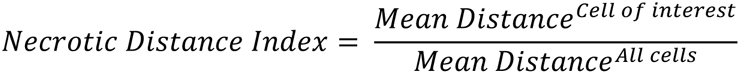

### Lesion neighbourhood and trajectory Analysis

The minimum distance between each non-necrotizing lesion to a necrotizing lesion was calculated in R using the pointDistance function from the raster package^44^. All cells from one lesion were measured with regard to another and the minimum function applied to the result identified the minimum distance between two lesions.

Data from immune cell densities, central preference and cell location index collated into a matrix and functions from the SCORPIUS package^45^ were used to produce the trajectory plot and trajectory heatmap.

### Statistical analysis

All statistical analyses were performed in GraphPad Prism 9.1.2. Single comparisons were performed using a Mann-Witney test. Multiple comparisons were performed using a Kruskal Wallis test with Dunns multiple comparisons test. Paired single comparisons were performed using a Wilcoxon matched-pairs signed rank test. A p-value < 0.05 was considered statistically significant for all tests. All plots were generated using GraphPad Prism or R.

## Acknowledgments

We acknowledge Clifton Barry III and Laura Via for their thoughtful discussion. This work was supported by US National Institutes of Health (U01AI166309) and the Natural Science Foundation of China (91942315). A.S. and J.E. were supported by Australian Postgraduate Awards.

## Author contributions

Conceptualization, A.S. and C.G.F.; Investigation, A.S., J.E., and T.F.; Data analysis: A.S., E.P., T.F. and C.G.F.; Resources, J.S.E, Q.Y., D.L. B., J.D.E., W.J.B., U.P., and X.C.; Writing – Original Draft, A.S. and C.G.F.; Writing – Review & Editing, All authors; Funding Acquisition, C.G.F., X.C., J.D.E., D.L.B. and E.P.; Supervision, C.G.F.

---

**Supplemental Table 1.**
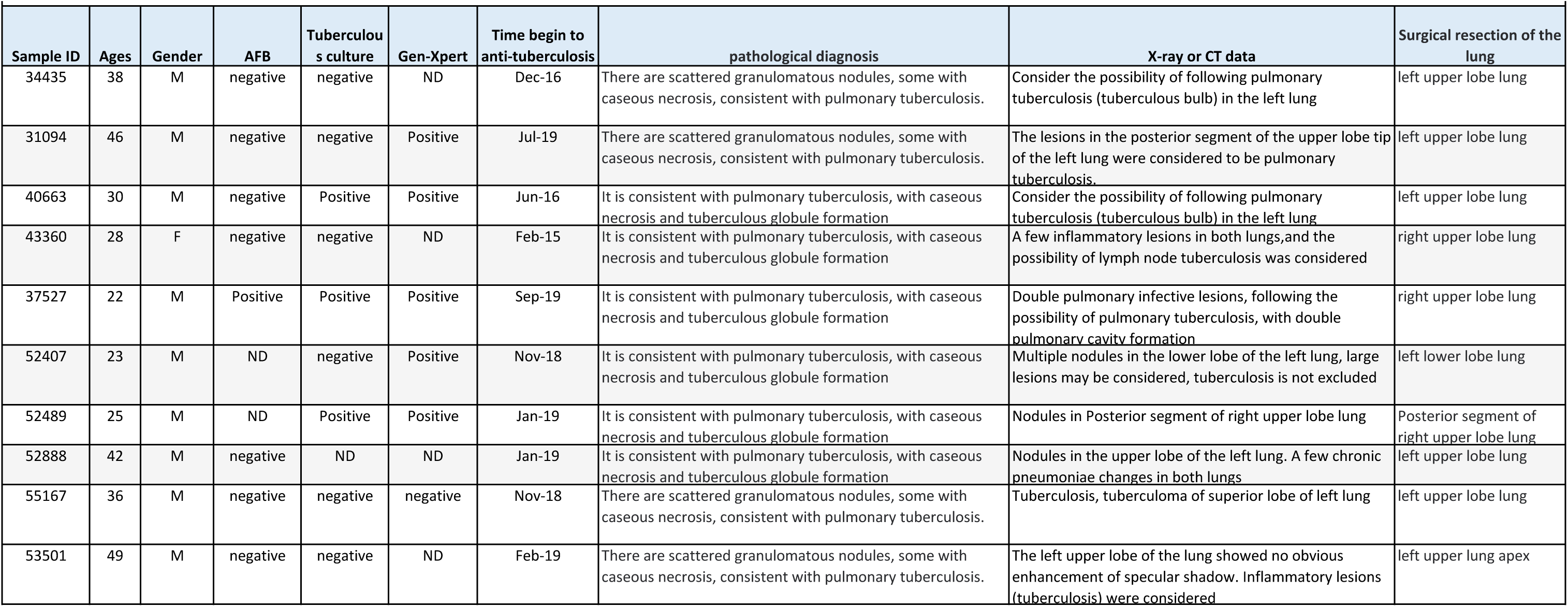
The cohort demographics and clinical information.

|  |  |  |  |  |  |  |  |  |  |
| --- | --- | --- | --- | --- | --- | --- | --- | --- | --- |
| Sample ID | Ages | Gender | AFB | Tuberculosis culture | Gen-Xpert | Time begin to anti-tuberculosis | pathological diagnosis | X-ray or CT data | Surgical resection of the lung |
| 34435 | 38 | M | negative | negative | ND | Dec-16 | There are scattered granulomatous nodules, some with caseous necrosis, consistent with pulmonary tuberculosis. | Consider the possibility of following pulmonary tuberculosis (tuberculous bulb) in the left lung | left upper lobe lung |
| 31094 | 46 | M | negative | negative | Positive | Jul-19 | There are scattered granulomatous nodules, some with caseous necrosis, consistent with pulmonary tuberculosis. | The lesions in the posterior segment of the upper lobe tip of the left lung were considered to be pulmonary tuberculosis. | left upper lobe lung |
| 40663 | 30 | M | negative | Positive | Positive | Jun-16 | It is consistent with pulmonary tuberculosis, with caseous necrosis and tuberculous globule formation | Consider the possibility of following pulmonary tuberculosis (tuberculous bulb) in the left lung. | left upper lobe lung |
| 43360 | 28 | F | negative | negative | ND | Feb-15 | It is consistent with pulmonary tuberculosis, with caseous necrosis and tuberculous globule formation | A few inflammatory lesions in both lungs, and the possibility of lymph node tuberculosis was considered | right upper lobe lung |
| 37527 | 22 | M | Positive | Positive | Positive | Sep-19 | It is consistent with pulmonary tuberculosis, with caseous necrosis and tuberculous globule formation | Double pulmonary infective lesions, following the possibility of pulmonary tuberculosis, with double pulmonary cavity formation | right upper lobe lung |
| 52407 | 23 | M | ND | negative | Positive | Nov-18 | It is consistent with pulmonary tuberculosis, with caseous necrosis and tuberculous globule formation | Multiple nodules in the lower lobe of the left lung, large lesions may be considered, tuberculosis is not excluded | left lower lobe lung |
| 52489 | 25 | M | ND | Positive | Positive | Jan-19 | It is consistent with pulmonary tuberculosis, with caseous necrosis and tuberculous globule formation | Nodules in Posterior segment of right upper lobe lung | Posterior segment of right upper lobe lung |
| 52888 | 42 | M | negative | ND | ND | Jan-19 | It is consistent with pulmonary tuberculosis, with caseous necrosis and tuberculous globule formation | Nodules in the upper lobe of the left lung. A few chronic pneumoniae changes in both lungs | left upper lobe lung |
| 55167 | 36 | M | negative | negative | negative | Nov-18 | There are scattered granulomatous nodules, some with caseous necrosis, consistent with pulmonary tuberculosis. | Tuberculosis, tuberculoma of superior lobe of left lung | left upper lobe lung |
| 53501 | 49 | M | negative | negative | ND | Feb-19 | There are scattered granulomatous nodules, some with caseous necrosis, consistent with pulmonary tuberculosis. | The left upper lobe of the lung showed no obvious enhancement of specular shadow. Inflammatory lesions (tuberculosis) were considered | left upper lung apex |

**Figure S1.**
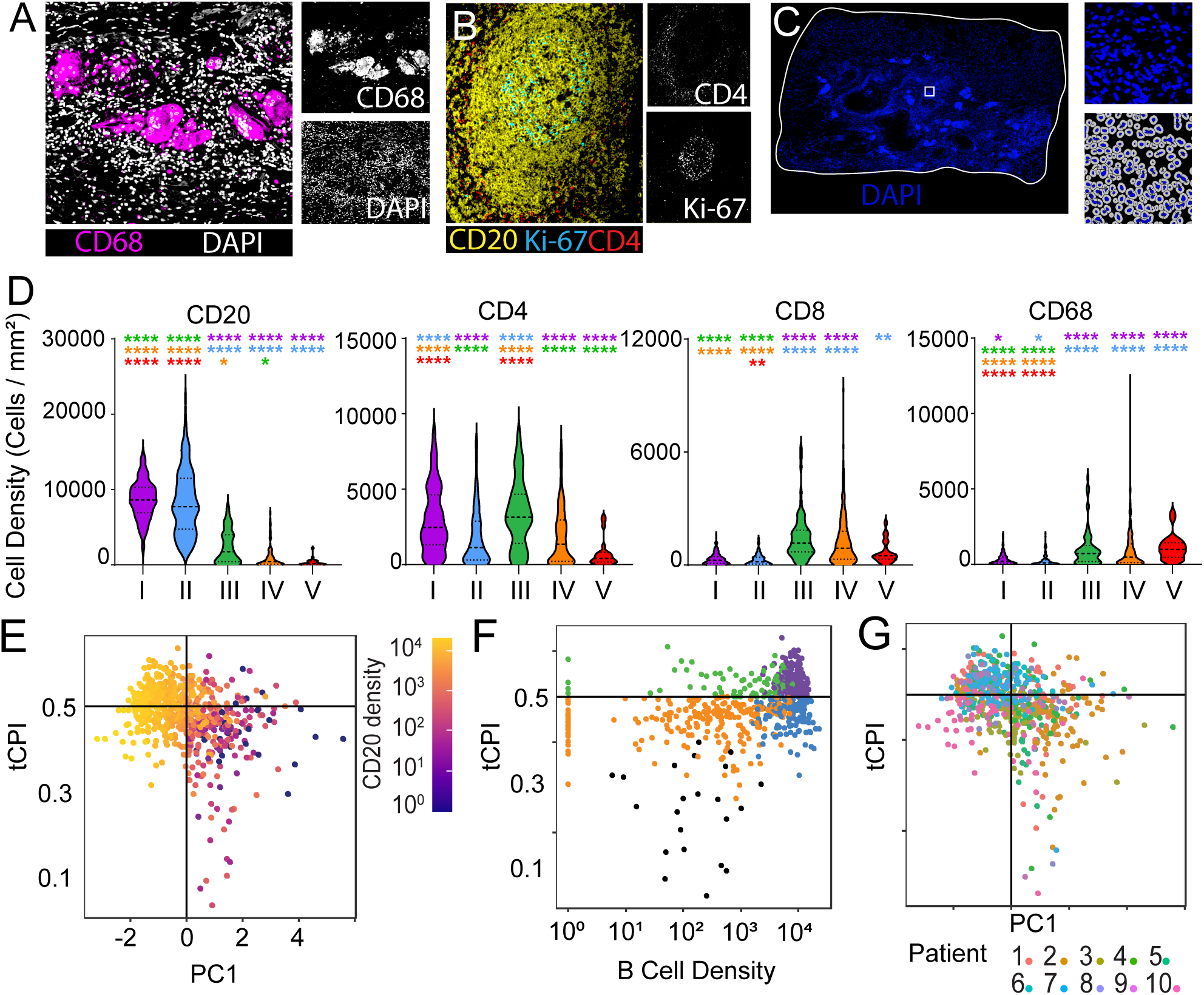
Lesion features in the M.tb-infected lung. **(A)** Representative image of Giant cells identified based on morphology and CD68 and DAPI staining. **(B)** Representative multiplex staining with CD20, CD4 and KI-67 showing a Germinal centre-like structure. **(C)** The same composite image of TB lung section as in Fig. 1A stained with DAPI with a representative view of segmented cells shown. **(D)** Density of major immune populations according to lesion type. The level of statistical significance is determined using a Kruskal-Wallis Test with Dunns Multiple Comparisons Test and an adjusted p-value less than 0.05 was considered statistically significant. * *P* < 0.05, ** *P* < 0.01, *** *P* < 0.001, **** *P* < 0.0001. **(E)** Comparison of PC1 against tCPI from Fig. 3E coloured according to density of CD20^+^ B cells. **(F)** Comparison of CD20^+^ B cell density against tCPI. Each symbol represents an individual lesion and lesions are coloured according to type. A value was of 1 was added to the density of all lesions to allow lesions with no CD20^+^ B cells to be displayed on the log scale. **(G)** Pairwise comparison of PC1 against tCPI. Each colour represents a lesion type.

**Figure S2.**
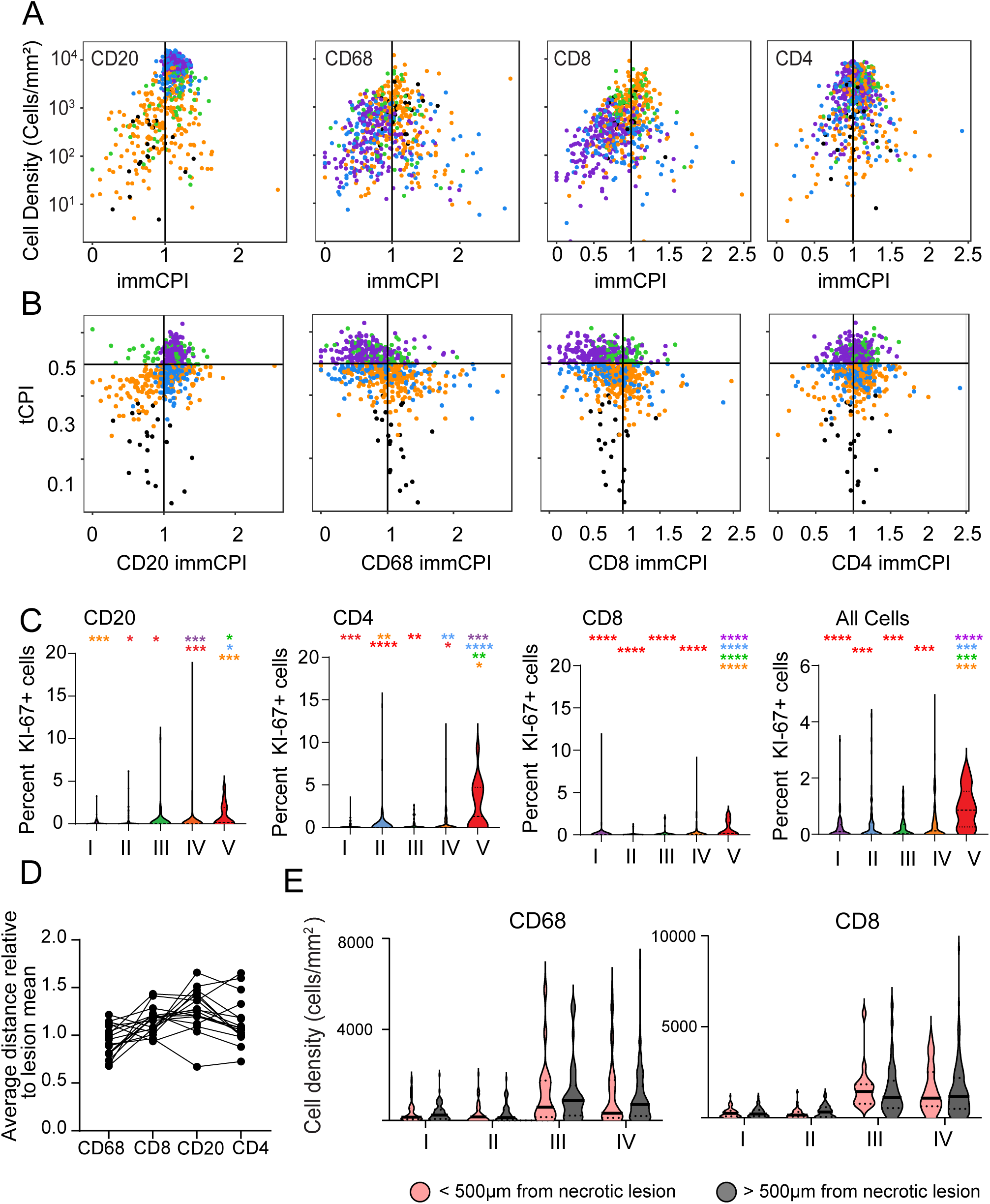
Spatial arrangement of cells in the M.tb-infected lung. **(A)** Pairwise Comparison of cell density versus immCPI for major immune populations. Each symbol represents an individual lesion and are coloured according to lesion type. **(B)** Comparison of tCPI against immCPI for major immune populations. Each symbol represents an individual lesion and are coloured according to lesion type. **(C)** Proportion of major cell populations positive for Ki-67. The level of statistical significance is determined using a Kruskal-Wallis Test with Dunns Multiple Comparisons Test and an adjusted p-value less than 0.05 was considered statistically significant. * *P* < 0.05, ** *P* < 0.01, *** *P* < 0.001, **** *P* < 0.0001. **(D)** Necrotizing Cell Index of major immune populations. Each symbol represents an individual necrotizing lesion and lines track data from the same lesion. **(E)** Density of CD8^+^ and CD68^+^ cells in lesions located greater or less than 500 µm from necrotizing lesions. The level of statistical significance is determined using a Mann-Whitney test and a p-value less than 0.05 was considered statistically significant. ** *P* < 0.01 and *** *P* < 0.001.

